# The Two Lives of Visual Working Memory: Evidence for Distinct Conscious and Unconscious Representations

**DOI:** 10.64898/2026.05.01.722131

**Authors:** Aleksandra Lipińska, Kinga Ciupińska, Renate Rutiku

**Affiliations:** Consciousness Lab, Institute of Psychology, Jagiellonian University, Krakow, Poland; Centre for Brain Research, Jagiellonian University, Krakow, Poland

**Keywords:** visual working memory, consciousness, perceptual awareness, conscious and unconscious processing, mental imagery

## Abstract

Visual working memory (vWM) is often linked to conscious experience and visual imagery, but it is typically described as a system that stores separate, independent items. These assumptions are difficult to reconcile, given the unified nature of conscious experience. Here, we test the hypothesis that vWM relies on at least two distinct representations: an underlying, unconscious memory trace and a consciously accessible, integrated representation.

A total of 216 participants performed a change-detection task, in which they rated their perceptual awareness of the memory display during the maintenance interval. Critically, we manipulated the statistical properties of the displays (average item size and size variability) to probe sensitivity to unified ensemble-level structure. Results revealed a dissociation between subjective and objective measures. Perceptual awareness increased for displays with larger, more variable items, whereas objective performance improved for displays with smaller, less variable items.

Despite this difference, subjective awareness still predicted performance, and even incorrect responses showed consistent biases rather than random guesses. Importantly, individual differences in imagery vividness (VVIQ) were selectively associated with subjective awareness and estimation bias, but not with objective correctness. These precision biases were further shaped by display statistics, suggesting that multiple representations can guide behavior.

Together, our findings support a reinterpretation of vWM performance in which task responses can draw on both unconscious and consciously accessible representations. One possible explanation for these behavioral patterns is that subjective experience reflects integrated, ensemble-like representations, while objective performance depends more strongly on item-specific information.

**Public significance statements:** Working memory allows us to temporarily hold and use information, and differences in this ability are closely linked to broader cognitive skills such as intelligence. This study shows that these differences may not depend only on how much information people can store, but also on how they experience it: some individuals appear to rely more on consciously accessible, image-like representations, especially when memory is uncertain or prone to error. By demonstrating that subjective experience and the vividness of imagery can shape behavior independently of objective accuracy, these findings suggest that how we use memory may be as important as how much we can store, with implications for understanding individual differences in cognition.

## 1. Introduction

Visual Working Memory (vWM) is a central function of cognition. It allows us to keep information active in our minds and use it flexibly for goal-directed tasks (Cowan, 2017). Yet our insight into exactly how vWM operates remains inconclusive. This study is focused on reconciling two prevailing yet conflicting assumptions about the basic properties of vWM. On the one hand, visual working memory performance is assumed to be a conscious process (Skóra & Wierzchoń, 2016). On the other hand, visual working memory capacity is typically estimated as if the to-be-remembered units of information (e.g., a set of objects or a display of colored squares) are stored independently (Luck & Vogel, 2013). There is much disagreement as to the definition of consciousness, but if there is one aspect that most accounts agree on, it is its unified nature (Bayne & Chalmers, 2003; Brook & Raymont, 2006; Seth & Bayne, 2022). Therefore, if visual working memory content is experienced consciously, its items cannot exist independently of one another. Below, we will elaborate on this point and propose that this paradox can be resolved by additionally assuming, as is proposed by the ‘conscious copy’ model (Jacobs & Silvanto, 2015), that there exists at least two different vWM representations, one unconscious and one conscious. From this, we derive and test the prediction that, if more than one representation exists, goal-directed tasks involving vWM content can be performed on any of them.

But first, to further explain the conflicting assumptions this study addresses, one should note that classical theories of vWM placed it firmly within the realm of conscious thought (Atkinson & Shiffrin, 1971). Presumably, this strong assumption was first made because most people believe they can consciously access the information while it is held in vWM. Many people would also describe this experience as perceptual, which is why a link between vWM and mental imagery has been prominent in the literature for some time (Baddeley & Andrade, 2000). In fact, the episodic buffer was specifically added to Baddeley’s working memory model to account for the conscious access into WM content (Baddeley, 2003).

At the same time, classic theories also assume that vWM performance in typical experimental tasks relies on a ‘slots’ model of change detection (Pashler, 1988; Cowan, 2001). According to this model, the bottleneck in vWM performance is due to a hard limit on the number of items that can be remembered because items are stored in separate preexisting ‘slots’ (Luck & Vogel, 1997). This practice of estimating vWM capacity as if the items in it are fully independent units persists to this day (Brady & Tenenbaum, 2013). For example, recent studies using the change detection paradigm continue to estimate vWM capacity based on accuracy as a function of set size, treating performance as a proxy for the number of stored items (e.g., Balaban et al., 2019; Zhao et al., 2023; Robinson et al., 2025). The assumption is that if a participant can retain X items in vWM but there are X+Y items in the to-be-remembered display, then the participant’s average performance should be approximately X/(X+Y) percent correct.

If one accepts the premise that conscious experience is unified, then it should be obvious that the ‘slots’ model is at odds with the claim that vWM is conscious. One or the other assumption (or both) cannot be strictly speaking correct. And indeed, evidence exists to call both assumptions into question. Studies have consistently shown that vWM performance is sensitive to the gist and ensemble properties of the memory displays, suggesting that the items are not maintained independently (for a review, see Brady, Konkle, & Alvarez, 2011). Haberman, Brady, and Alvarez (2015) even demonstrated the existence of more than one level of ensemble representations in hierarchical displays. This would fit well with the conscious vWM account. The content of vWM also seems to be preferentially chunked (Orbán et al., 2008; Orhan & Jacobs, 2014; Nassar, Helmers & Frank, 2018), which, again, is much more similar to the way humans consciously perceive visual information, rather than to a purely independent ‘slots’ model.

On the other hand, the trouble with the conscious account of vWM performance is that, despite our impression of conscious access to WM content while maintaining it, we are generally quite bad at using this information in subsequent WM tasks, e.g., comparing the memory to a test item. In a typical vWM task, it is as if, once faced with the test display, we are baffled at how difficult it is to detect the changed item. This suggests that we may not have as much conscious access to the processes associated with vWM as we believe. This interpretation is supported by a series of studies demonstrating that vWM can operate unconsciously to some extent (Soto, Mäntylä, & Silvanto, 2011; Trübutschek et al., 2017; Bergström & Eriksson, 2018; Franco-Martínez et al., 2026). Importantly for the purposes of this study, it should be noted that if some kind of unconscious vWM capacity exists, it would be much more likely that this capacity can operate on the basis of separate elements, akin to the ‘slots’ model of change detection. The conscious account of vWM performance also faces another problem. As mentioned above, a strong link between vWM retention and mental imagery is often assumed (Keogh & Pearson, 2011; Bates et al., 2024). However, people with aphantasia who report no mental imagery during the retention period still perform just as well on vWM tasks as those who report having a mental image of the memory display (Keogh & Pearson, 2018). This consistent finding has raised criticism against the standard vWM models, or at least their assumption of conscious access (Pearson, 2019).

The final problem with the conscious account of vWM is that it relies almost entirely on face validity, as it aligns well with most people’s spontaneous introspection about their vWM performance. If one does not take confidence reports into account (e.g., Rademaker et al., 2012; Vandenbroucke et al., 2014; Conte et al., 2023), this assumption has not been rigorously tested (although initial steps have been taken by, e.g., Baddeley & Andrade, 2000; Mikels et al., 2008; Forsberg, Blume & Cowan, 2021). We lack detailed data on how people actually consciously perceive their vWM content, particularly its perceptual nature. Moreover, most studies that have incorporated subjective perceptual ratings in their task design have applied them after the objective vWM task, not during vWM retention, i.e., while people are actively maintaining the task-relevant vWM representations in their mind. If we wanted to better understand whether vWM performance operates on conscious memory representations and whether this has any functional implications, we should ask people about perception during memory maintenance.

Yet, despite the lack of empirical data, it is irrefutable that most people believe they have *some* conscious experience of their vWM content. This fundamental realization, together with a well-reasoned analysis of the vWM literature, led Jacobs and Silvato (2015) to propose the ‘conscious copy’ model of vWM introspection. The model proposes that the maintenance of vWM content is fundamentally unconscious, but introspection of that content requires the creation of a conscious copy, i.e., a new representation. The model further assumes that these two representations are independent, and whereas the vWM representation can be feature-specific, the conscious representation includes bound objects in the same way as they are perceived in episodic memory. For completeness (yet not directly relevant for the present study), the final assumption is that the two representations interact differently with incoming sensory information (for further development of the model, see also Jacob, Jacobs & Silvanto, 2015).

In light of the previously described problems with two prevalent assumptions about vWM, the ‘conscious copy’ model seems to offer an appealing solution. But it also opens up the possibility of an even more radical reinterpretation of vWM performance. Namely, it suggests that any questions regarding vWM content (not only introspection) can be answered based on the unconscious vWM representation as well as the ‘conscious copy’ of the memory trace. To put it differently, the creation of the ‘conscious copy’ may be a habitual process for many people, regardless of whether they are explicitly required to introspect about the vWM content. In situations where the vWM task is demanding, and the capacity of the unconscious store is exceeded (as is the case in many typical experimental tasks), it is perhaps even more likely that the task is performed based on the ‘conscious copy’ representation. For example, if the task is change detection, and on a given trial the unconscious vWM store fails to support correct identification of the changed item (regardless of whether the fault was with encoding, maintenance, or decoding of the correct information), then the participant may still resort to the ‘conscious copy’ representation to discern a best guess.

The present study tests the assumption that the contents of vWM are reflected in multiple representations. We build on the widely accepted premise that conscious experience is unified (Bayne & Chalmers, 2003; Bayne, 2010). If conscious and unconscious vWM representations differ in their level of integration, then subjective and objective vWM-related responses should also differ in their sensitivity to the ensemble properties of the memory displays. To test this, we systematically manipulate ensemble properties of vWM items and assess whether subjective experience reflects integrated, ensemble-like representations, as predicted by a unified conscious percept. Critically, we compare the influence of these ensemble properties on subjective ratings versus objective performance. In addition, we also examine whether these putative conscious and unconscious representations operate independently or whether task performance can draw on both kinds of representations. Specifically, we evaluate whether the precision of vWM responses across correct and incorrect trials is best characterized by a mixture of representational formats.

## 2. Methods

### 2.1 Participants

The experiment was realized as part of the EU COST Action CA18106 (www.neuralarchcon.org), at the data collection site in Kraków (Consciousness Lab, Jagiellonian University, Kraków, Poland). The broader consortium project encompasses an extensive array of behavioral tests, MRI sequences, EEG experiments, and DNA probing of neurotypical participants. The study at hand focuses on the behavioral results of one visual working memory task of the larger test battery.

The final sample after preprocessing (see Section 2.5 “Preprocessing”) consisted of 216 participants (79 male) out of 253 participants who completed the visual working memory task. Participants were between 18 and 39.1 years old (*M* = 23.62, *SD* = 4.09). All individuals reported good physical health at the time of testing, had normal or corrected-to-normal visual acuity, and normal hearing. Exclusion criteria consisted of any prior history of brain injury or neurosurgical intervention, diagnosed structural brain abnormalities, and the use of neuropharmacological agents or other medications with potential central nervous system effects. All participants met standard MRI safety requirements and provided written informed consent before enrollment. The study was conducted in accordance with the Declaration of Helsinki and received approval from the Ethics Committee of the Institute of Psychology, Jagiellonian University.

### 2.2 Stimuli

The working memory displays consisted of four striped circles presented in the four quadrants of the visual field (Figure 1A; see SI.1 for additional examples). Circle size was the only task-relevant feature. Stripe orientation was fully randomized, and stimulus positions were varied across trials to minimize visual habituation and discourage visually driven strategies (Manassi, Murai & Whitney, 2023).

**Figure 1.**
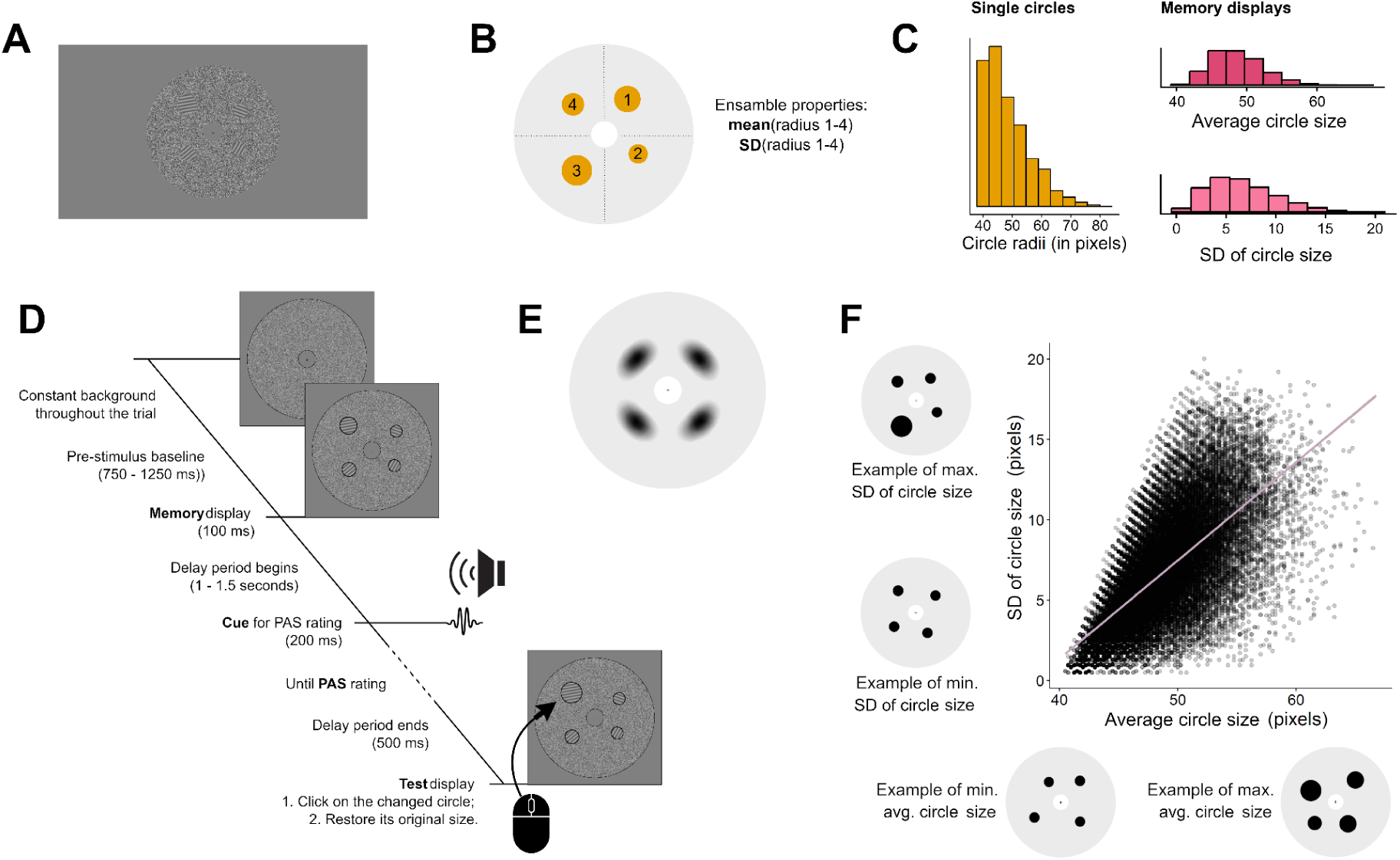
Paradigm illustrations. **A**. An example of a memory display, unaltered. Note that the circles were easy to see on the monitor during the experiment, even though they appear low-contrast in the illustration. **B**. All memory displays consisted of four circles of varying sizes, placed in the four quadrants of the background annulus. Two ensemble properties were derived from the set of four circles: the average circle size and the standard deviation of circle size. **C**. (left) Distribution of the size of the individual circles across the dataset. The size is measured as the radius of a circle in pixels. (right) Distributions of the average circle size and the standard deviation of circle size across all the memory displays in the dataset. These distributions are depicted in more detail in Figures 3A and 3E. **D**. A graphical summary of the trial structure. The stimulus edges have been highlighted for better visibility. **E**. A heatmap of all the circles in the dataset. **F**. All combinations of average circle size and standard deviation of circle size. Each dot represents one trial in the dataset. Examples of the extreme displays for both ensemble properties are depicted along the axes.

Circle sizes were drawn from an exponential distribution ranging from 0.7 to 1.4° of visual angle (43 discrete values; Figure 1C). On each trial, four sizes were sampled with replacement and assigned to the four circles. The use of an exponential distribution, inspired by Groen and colleagues (2012), ensured a clearer dissociation between the probability of individual item sizes and the mean size of the four-item display.

Each circle was assigned to a fixed quadrant, but its exact position within that quadrant varied across trials. Positioning followed a stochastic procedure. First, circle size was determined as described above. Next, eccentricity (distance from fixation) was sampled uniformly between 3° and 4° of visual angle. The polar angle was then drawn from a truncated normal distribution centered on the midpoint of the assigned quadrant. Specifically, angular ranges were 10–80° (mode 45°) for the upper-right quadrant, 100–170° (mode 135°) for the upper-left quadrant, 190–260° (mode 225°) for the lower-left quadrant, and 280–350° (mode 315°) for the lower-right quadrant. This approach increased the likelihood of positions near quadrant centers while maintaining variability across trials. Cardinal meridians (0°, 90°, 180°, 270°) were excluded to avoid placement directly along the horizontal and vertical axes.

After determining polar coordinates, stimuli were provisionally rendered and checked for spatial overlap. A buffer zone of 1° of visual angle was added around each circle, and any overlap (including buffer zones) triggered a full resampling of positions for that trial. This process was repeated until all four circles were non-overlapping. Figure 1E shows a heat map of possible stimulus locations.

The orientation of the stripes ranged between 10 and 360 degrees (in steps of 10 degrees). On every trial, four orientations were randomly chosen from this set of 36 angles without replacement and assigned to the four circles of the working memory display. The orientations remained unchanged in the test display.

The task-relevant set of circles was superimposed on a larger background annulus consisting of white noise. This noise annulus was used to prevent afterimages of the striped circles. The radius of the inner border of this noise annulus was 1 degree of visual angle, and the radius of the outer border was 7 degrees of visual angle. The white noise was centered on the luminance level of the uniform gray background with a contrast variance of 60%. The luminance of the gray background was approximately 46 candela per square meter (cd/m^2^), and the average luminance of the noise annulus was approximately 55 cd/m^2^. The average luminance of the striped circles was approximately 50 cd/m^2^. The size of the central fixation cross was approximately 0.3 degrees of visual angle.

Taken together, the working memory displays were designed to control the task-relevant dimension of size. This encompasses both the sizes of the individual circles and their average size and variance (see below). On the other hand, other visual aspects of the stimuli, specifically orientation and location, were designed to vary across trials to reduce their systematic impact on the visual working memory performance and therefore allow for more generalizable results. The target of the working memory task was a change in the size of one of the four circles on each trial. The following three target-related factors were controlled for. First, the target could appear equiprobably in one of the four quadrants of the screen. Second, the target’s relative size compared to the other three circles was selected such that the target was equally likely to be the smallest circle, the second smallest circle, the second largest circle, or the largest circle in the display. This was achieved by ordering the randomly drawn circle sizes on every trial and assigning the appropriate one to the target. Third, the target was equally likely to decrease or increase in size compared to its original radius. The size change was always −20% or +20% of the target’s original size. Note, however, that these target-related factors are not of primary interest in the present study, and their influence on objective vWM performance is only covered in SI.3 for completeness.

### 2.3 Experimental procedure

The trial structure is depicted in Figure 1D. Example video recordings of the procedure are available for download at https://osf.io/6q43x/overview. Throughout the experiment, a central fixation cross and a noise annulus were displayed on the gray background. At the beginning of each trial, the white noise of the annulus was resampled and redrawn on the screen. The white noise remained static for the remainder of the trial. Participants were instructed to maintain fixation on the small central cross throughout the trial (except when giving the working memory response; see below) to improve performance. After 750–1250 ms, the working memory display consisting of four circles was superimposed on the noise annulus for 100 ms. Participants were instructed to memorize the sizes of the four circles as best they could. After a 1–1.5-second delay, a simple beep signaled to participants that they should now rate how clearly they are experiencing the memory display in their minds. Ratings were given on a four-point scale similar to the widely used Perceptual Awareness Scale (PAS; Ramsøy & Overgaard, 2004). Participants were given a detailed verbal explanation at the beginning of the experiment of what it means to have a conscious experience of vWM content, and that it is okay to have no conscious experience. Emphasis was put on the subjective nature of this experience, and that the following rating options should be used:

1. – No Experience. The participant has no conscious experience of the contents of working memory during maintenance.
2. – Faint experience. The participant is faintly experiencing the memory of the stimulus display during maintenance, but the experience is very vague, blurry, or partial.
3. – Almost Clear Experience. The participant experiences the memory of the stimulus display quite clearly during maintenance.
4. – Clear Experience. The participant experiences the memory of the stimulus display clearly during maintenance, almost as if it is still lingering on the screen.

For simplicity, we will continue to refer to this scale as PAS throughout the rest of the work. 500 ms after a PAS rating was given on the keyboard, the test display was superimposed on the noise annulus. In every test display, one of the four circles was smaller or larger than its original size in the memory display, while the size of the other three circles remained unchanged. Participants had to first click on the circle, which they thought had changed size relative to the memory display. Second, while holding down the left mouse button and moving the cursor, they had to restore the selected target item to its original size. (Note that the mouse cursor was only visible while the test display was active and participants gave the working memory response.) Once the participants were satisfied with their answer, they could start the next trial by pressing the space bar.

In the beginning, participants received 5 practice trials to get acquainted with the task and have a chance to ask questions. During the main portion of the experiment, two short self-paced breaks were scheduled after every 63 trials. The intended length of the experiment was a minimum of 192 trials or ca. 30 minutes of task duration, but due to the demands of large-scale data collection, not all participants completed all trials; some completed fewer trials than planned, while others completed a surplus if recording time remained. The number of available trials for analysis is detailed at the end of the next section.

Participants were seated in a dimly lit room, approximately 60 cm away from the screen (23.8-in. HP monitor; NVIDIA GeForce RTX 2070 SUPER graphics card; 1920 × 1080 screen resolution; 60 Hz refresh rate). The PAS ratings were given with the left hand on the computer keyboard (number keys 1, 2, 3, and 4). The beep that cued the PAS rating was played through earphones. It was a C5 chord (523, 1046, and 1569 Hz) with a 200 ms duration and 10 ms up-down ramps. The working memory task was performed with a standard computer mouse.

### 2.4 Vividness of Visual Imagery Questionnaire

Participants’ ability to generate mental images was assessed using the 32-item *Vividness of Visual Imagery Questionnaire-2* (VVIQ-2; Marks, 1995). The VVIQ is a self-report measure designed to capture individual differences in the subjective clarity and vividness of voluntary mental imagery. It has been widely used and validated as a measure of imagery ability in both cognitive and consciousness research (Marks, 1973; McKelvie, 1995). A Polish translation of the questionnaire composed by the project team was used in the present study.

Participants were instructed to generate mental images of a series of described scenarios (e.g., “a sun rising above the horizon into a hazy sky”) and rate the vividness of each image on a 5-point Likert scale, ranging from 0 (“No image at all, you only know that you are thinking of the object”) to 4 (“Perfectly clear and vivid as real life”). Following previous approaches (Zeng et al., 2022; Kvamme et al., 2026), summed VVIQ scores were computed across all items and subsequently normalized to the unit interval, with higher values indicating greater subjective vividness of visual imagery.

### 2.5 Preprocessing

Participants’ data were removed from the subsequent analyses if **i)** more than 20% of trials were skipped without choosing a target or adjusting its size (14 participants removed); **ii)** the average precision error of the target’s size adjustment was very high (mean plus 3xSD; 2 participants removed); **iii)** the same PAS rating was chosen on all trials (3 participants removed); **iv)** the average correctness was 30% or less (11 participants removed); **v)** fewer than 100 trials were left after preprocessing (7 participants removed).

Individual trials were removed from subsequent analysis if **vi)** the PAS rating was not clear, i.e., more than one button was pressed at once (<0.1% of data); **vii)** the trial was skipped without choosing a target or adjusting its size (2% of data); **viii)** the size adjustment of the target took a very long time (mean plus 3xSD; 2.9% of data); **ix)** the standard deviation of item sizes on the final display, after responses were given, is very high (mean plus 3xSD; 0.5% of data); **x)** the selected target circle’s size was adjusted more than 4 times before moving on to the next trial (4% of data).

After preprocessing, the final sample comprised 216 participants (85.4% of the raw sample). The average number of remaining trials per participant was 177 (*SD* = 21, *range* = 117–252 trials).

### 2.6 Statistical analysis

The central aim of this study was to characterize the influence of task-related ensemble properties on subjective PAS ratings and objective vWM performance. Two ensemble properties are analyzed for that purpose – the average circle size and the standard deviation of circle size on each trial (where the individual display on each trial always consists of four circles; Figure 1B). Figures 3A and 3E depict the distribution of these two ensemble properties for the entire dataset. Before statistical analysis, however, both distributions were further divided into quantiles for each participant separately and coded as ordinal variables. This helps increase the coherence and robustness of the analyses, since the exact distribution of the ensemble properties varied slightly across participants, and some of the observed result patterns for objective vWM performance were nonlinear.

The overall analysis strategy was to fit two statistical models – a null model and an alternative model – for each analysis and to assess the significance of a predictor by comparing the models using an appropriate likelihood ratio test (reported as a chi-squared statistic, *Χ*^2^) and qualitatively evaluating the change in the Akaike Information Criterion (AIC^change^ = AIC of the alternative model - AIC of the null model). Since this experiment was limited in the number of trials each participant performed, interactions between predictors are analyzed sparingly. Instead, most alternative models include only one fixed-effect predictor of interest, and the null models include only the intercept (all exceptions to this pattern are noted in the Results). All fitted models also included a random-effect term for participant.

The effects of the two ensemble properties on PAS ratings were assessed using Cumulative Link Mixed Models with a logit link function, and likelihood ratio tests were performed using the Nagelkerke pseudo R-squared method. The effects of the two ensemble properties on %correct were assessed using Binomial Generalized Linear Mixed Models with a logit link function, and likelihood ratio tests were performed using the standard ANOVA method. The effects of the two ensemble properties on precision were assessed using Linear Mixed Models, and likelihood ratio tests were again performed using the standard ANOVA method. All follow-up pairwise comparisons between factor levels were conducted with paired t-tests. Pairwise comparisons were only conducted if the overall model comparison was significant.

The analysis of the influence of PAS ratings on objective vWM performance followed the same rationale as for the two ensemble properties, and PAS was coded as an ordinal variable. The relationship between VVIQ scores and the performance metrics of this experiment was assessed via Pearson correlations. Differences between distributions were assessed via asymptotic two-sample Kolmogorov-Smirnov tests.

## 3. Results

### 3.1 Subjective and objective measures of working memory

#### 3.1.1 PAS ratings during vWM maintenance

In contrast to previous studies, participants rated their perceptual awareness of vWM content while it was actively maintained. At the time of the rating, participants did not yet know how the stimulus display would change and which of the four circles would be the target. The PAS ratings are therefore unpolluted by the objective task performance and should more closely reflect people’s vWM experience.

The average PAS rating of the ongoing vWM content was 2.1 (sd=.56; range=1.01-3.9; Figure 2A). Figure 2D shows the proportions of the four response options. The most commonly chosen rating was PAS 2 (corresponding to a faint experience). Still, most people used more than two different response options over the course of the experiment (mean=3.28 out of four response options). It can therefore be concluded that PAS ratings exhibit sufficient range in this experiment and reflect the trial-to-trial variance in the online perception of the vWM content.

**Figure 2.**
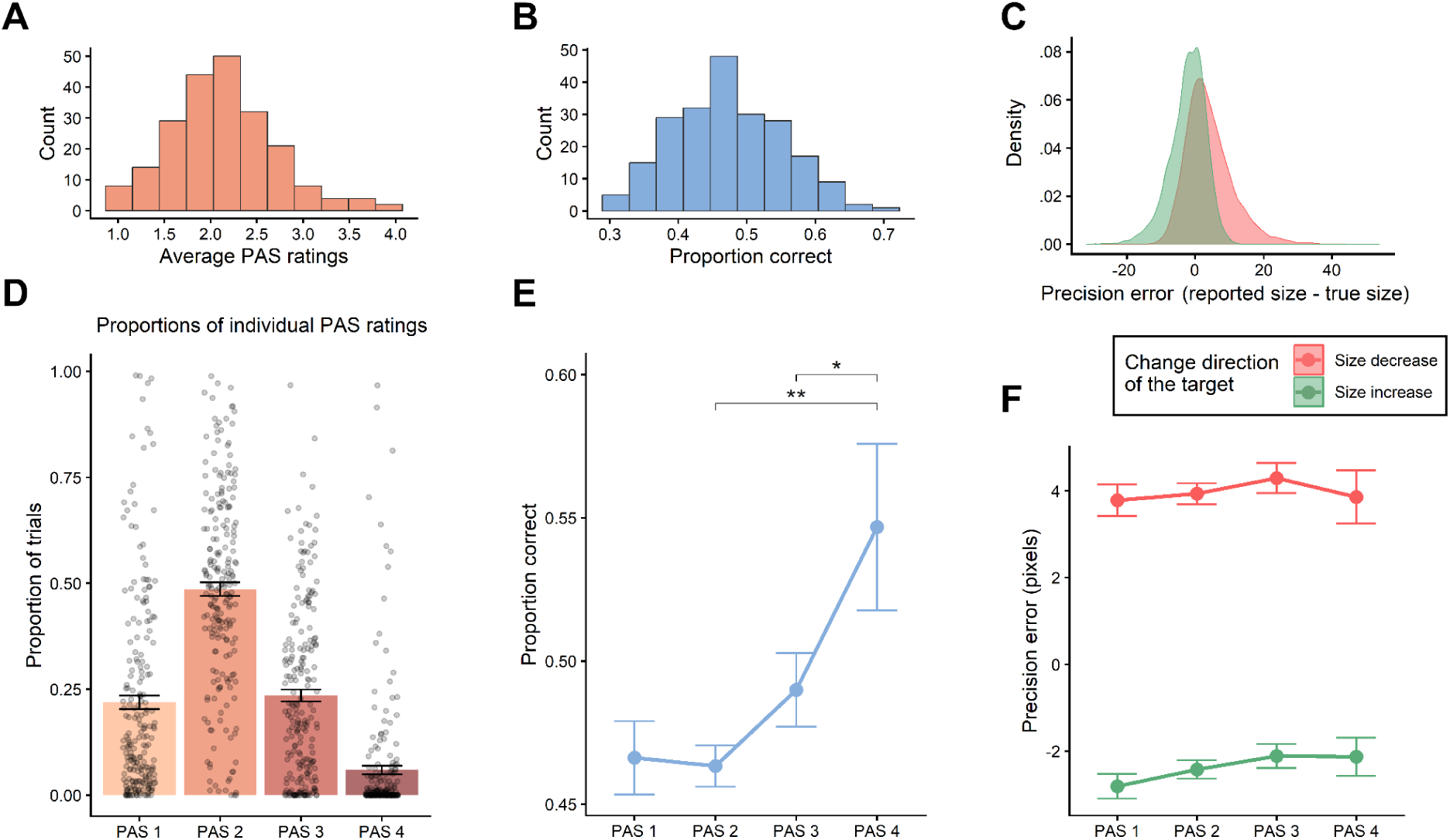
Descriptive statistics for subjective and objective vWM measures. **A**. Distribution of average PAS ratings across participants. **B**. Distribution of average objective performance (%correct) across participants. **C**. Distribution of precision error, i.e., how closely the size adjustment reflected the true original size of the target circle, across participants. Precision error is shown separately for increases and decreases in the target’s size. **D**. Proportions of how often the individual PAS ratings were chosen by participants. Each dot represents one participant per PAS rating. Error bars depict standard error estimates. **E**. Average proportion of correct trials, depending on which PAS rating preceded the target selection. Error bars depict standard error. Significance bars depict pairwise comparisons between PAS ratings with a significant t-statistic. Note that the number of participants available for each pairwise comparison differed because not all participants used all four PAS ratings. **F**. Average precision error, depending on which PAS rating preceded the target’s size adjustment. Precision error is shown separately for increases and decreases in the target’s size. Error bars depict standard error.

#### 3.1.2 Correctness

Correctness refers to whether or not the correct target, i.e., the circle that indeed changed size, was selected on a given trial. On average, participants chose the correct target on 47% of trials (sd=8%, range=31-70%; Figure 2B). This means that the task was quite difficult, but performance remained consistently above chance.

#### 3.1.3 Precision of correct responses

In addition to identifying the target circle, participants also had to estimate its original size. Precision refers to how close this estimate is to the true original size of the selected circle. Since this measure is only meaningful as an objective index of vWM performance if the correct target circle was selected in the first place, the following analyses focus solely on the correct trials for now. Furthermore, since precision was strongly influenced by the change direction of the target circle’s size (see SI.3.2), this factor is always included as a covariate.

The average objective precision of vWM performance was 1.1 pixels (sd=1.9 pixels; range=-5.5 – 9.2 pixels). Figure 2C shows the distribution of precision across the entire dataset, divided by change direction. There are two general observations regarding the distribution of precision. First, participants tend to overcorrect the target circle’s size. When the size decreased, the responses tended to overestimate the original circle size by an average of 4 pixels. When the size increased, the responses tended to underestimate the original circle size by an average of −2.2 pixels. Second, despite the overcorrection also being negative, participants generally still showed a slight preference for overestimation over underestimation.

#### 3.1.4 Is objective vWM performance predicted by PAS ratings?

If higher PAS ratings lead to relatively more correct target selections, this would indicate that both subjective and objective vWM performances are at least partly informed by the same representations. And indeed, there was a significant positive effect of PAS ratings on correctness (*Χ*^2^(3) = 32.9, p < .001; AIC^change^ = −27; Figure 2E). Participants tended to be systematically more correct when the preceding PAS rating was 4 compared to when the preceding PAS rating was 1, 2, or 3 (all t < −2.01, all p < .047). The other pairwise comparisons of PAS ratings were not significant (all t-values < −1.7, all p-values> .094).

The precision of target adjustment was not systematically predicted by PAS ratings (*Χ*^2^(6) = 4.7, p = .58; AIC^change^ = 7.3), nor was there a significant interaction between PAS ratings and change direction (*Χ*^2^(3) = 0.6, p = .89; AIC^change^ = 5.4). The change direction itself was, of course, a highly significant predictor of precision (*Χ*^2^(4) = 628.1, p<.001; AIC^change^ = −620.1). The average precision per condition is depicted in Figure 2F.

### 3.2 Ensamble properties affect subjective and objective working memory differently

Two ensemble properties are used to characterize the heterogeneity of the displays across trials. Average circle size denotes the mean of the four circles’ radii on every trial. The standard deviation of circle size denotes the standard deviation of the four circles’ radii on every trial. To more coherently represent the ensemble properties of the stimulus displays across participants and to better grasp nonlinear patterns, we divided both statistics into quantiles and evaluated the effects of these factors on the subjective and objective vWM measures separately. The reason for separate analysis is that the average item size and the standard deviation of item size are correlated (r = .67, p < .001; Figure 1F). Therefore, the point here is not to delineate their relative contribution, but to describe an overall pattern of results which can be compared between subjective and objective vWM measures.

#### 3.2.1 PAS ratings

Both ensemble properties had a significant effect on the PAS ratings (*Χ*^2^(4) = 281.6, p < .001, AIC^change^ = −273.59, and *Χ*^2^(4) = 479.2, p < .001, AIC^change^ = −471.23 for average circle size and the standard deviation of circle size, respectively). Table SI.2.1 lists the test statistics for a more detailed overview of these effects. As shown in this table, both linear and quadratic trends were significant for the two ensemble properties. Higher PAS ratings tended to occur on trials that had a larger average circle size (see Figure 3B). Specifically, quantiles 3, 4, and 5 yielded systematically higher PAS ratings than quantile 1 (all t < −3.4, all p < .001). Quantile 2 was not significantly different from quantile 1 (t(215) = −.5, p = .65). Higher PAS ratings also tended to occur on trials with a higher standard deviation in circle size (see Figure 3F). Specifically, quantiles 4 and 5 yielded systematically higher PAS ratings than quantile 1 (t(215) = −6, p < .001; t(215) = −10.7, p < .001, respectively). Quantiles 3 and 2 were not significantly different from quantile 1 (t(215) = −1.3, p = .02; t(215) = 0.8, p = .4, respectively).

**Figure 3.**
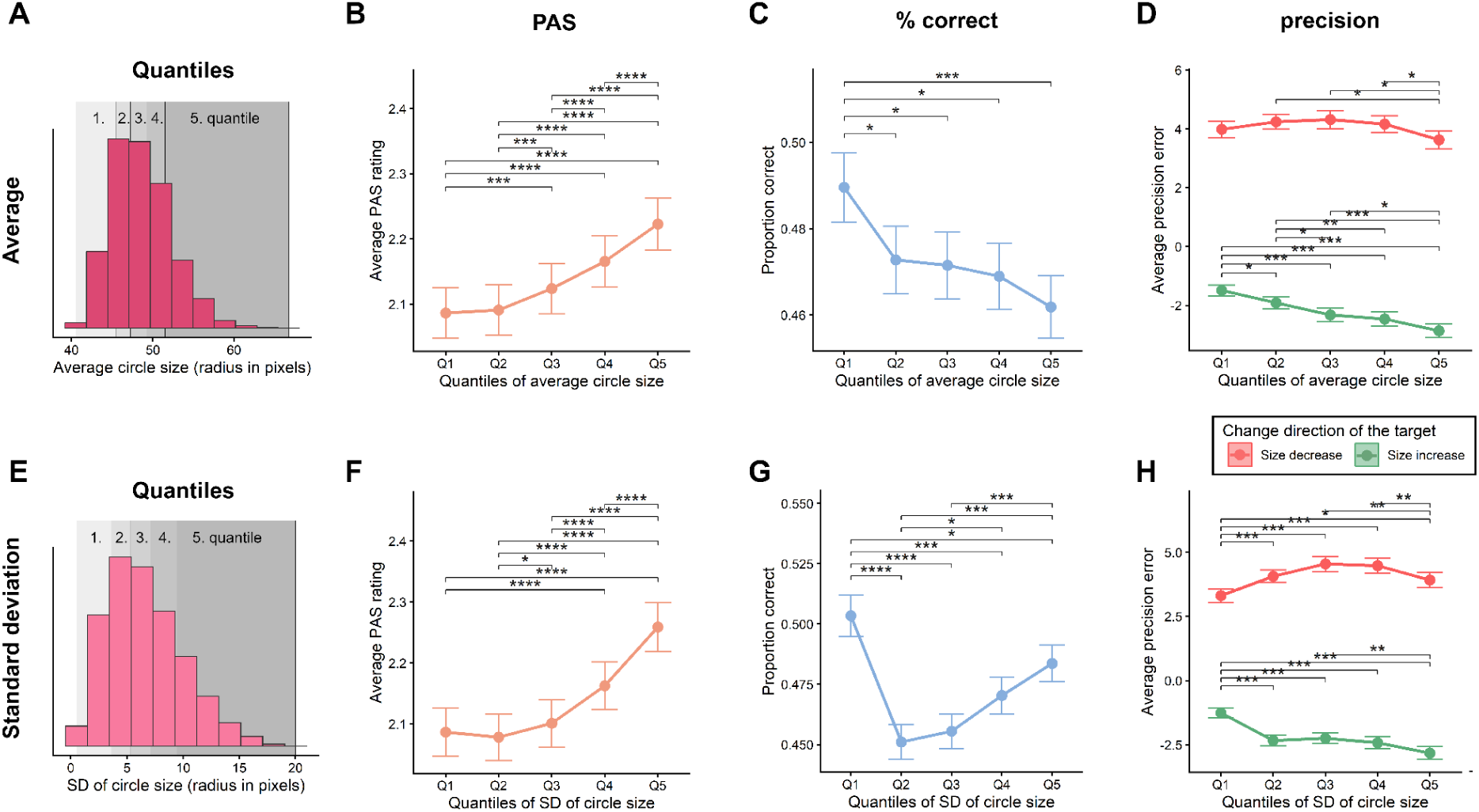
Effects of ensemble properties. **A & E**. Distribution of the average circle size and standard deviation of circle size across all the memory displays in the dataset. For analysis, the distributions were divided into quantiles on a participant-by-participant basis. The shaded areas illustrate approximately where the quantile borders were. **B**. Effect of average circle size quantiles on PAS ratings. **C**. Effect of average circle size quantiles on the proportion of correct responses. **D**. Effect of average circle size quantiles on precision, after accounting for the change direction of the target. **F**. Effect of SD of size quantiles on PAS ratings. **G**. Effect of SD of size quantiles on the proportion of correct responses. **H**. Effect of SD of circle size quantiles on precision, after accounting for the change direction of the target. In all the above plots, error bars depict standard errors, and significance bars depict pairwise comparisons between quantiles with a significant t-statistic.

#### 3.2.2 Correctness

Both ensemble properties also had a significant effect on the likelihood of choosing the correct target, but the pattern of results was not comparable to that of PAS ratings (see Figures 3C and 3G). A smaller average circle size tended to lead to more correct target selections (*Χ*^2^(4) = 12.8, p = .012; AIC^change^ = −15.1). Only the linear trend in the alternative model was significant (see table SI.2.2). The effect of the standard deviation of circle size was U-shaped (*Χ*^2^(4) = 56.4, p < .001; AIC^change^ = −48.5), and both the quadratic and cubic trends in the alternative model were significant. Quantile 1 yielded the highest proportion of correct trials, even compared to quantile 5 (t(215) = 2.05, p = .041), most likely due to a strong pop-out effect of the target in the test display. Quantiles 2 and 3 yielded the lowest proportions of correct trials, while correctness gradually increased again for quantiles 4 and 5.

#### 3.2.3 Precision

The average circle size had a significant effect on precision (*Χ*^2^(8) = 22.8, p = .004; AIC^change^ = −7), with no interaction with change direction (*Χ*^2^(4) = 7.8, p = .1; AIC^change^ = 0). Looking at the results for both change directions separately is nonetheless informative. Figure 3D shows that participants tended to overestimate the original circle size when the target’s size had decreased in the test display, with no clear effect of average circle size on the extent of overestimation. Conversely, participants tended to underestimate the original circle size when the target’s size had increased in the test display, and the average circle size had a clear effect on the extent of underestimation. Target adjustments tended to be more precise for displays with smaller circles and became increasingly biased towards underestimation as average circle size increased. The effect of the standard deviation of circle size was also significant (*Χ*^2^(8) = 38; AIC^change^ = −22), with a significant interaction with change direction (*Χ*^2^(4) = 30.9, p < .001; AIC^change^ = −23). The linear and quadratic terms were significant for the main effect and the interaction. The precision error tended to increase for displays with more diverse circle sizes, and this increase was again akin to a systematic overcorrection of the original target size, depending on the change direction (see Figure 3H).

Taken together, these results are akin to a mixture of the correctness and PAS result patterns. On the one hand, participants are more accurate for a smaller average circle size and less variance in the displays. On the other hand, responses do not become less accurate as average circle size and standard deviation increase; they become more biased towards overcorrection, indicating greater interaction among the display items.

### 3.3 Vividness of visual imagery and working memory experience

The above-described pattern of results suggests that the ensemble properties of the displays affect subjective vWM experience and objective vWM performance in different ways, but with some evidence of overlap as well. The findings are therefore only partly in line with the ‘conscious copy’ account of vWM, according to which the conscious experience of working memory content relies on an entirely separate representation. Further evidence supporting this conclusion comes from participants’ self-reported visual imagery vividness (assessed via VVIQ) and its relation to the vMW measures in this experiment.

The correlation between participants’ total VVIQ scores and average PAS ratings was significant (r = .26, t(210) = 3.8, p < .001; Figure 4B), indicating that people with higher VVIQ scores tended to give higher PAS ratings on average. The correlation between participants’ total VVIQ scores and average correctness was clearly not significant (r = .02, t(210) < 1; Figure 4C). The results for precision were a bit more curious, however. When looking at correct trials, VVIQ scores were not significantly associated with average precision, after accounting for the effect of change direction (*Χ*^2^(2) = 2.7, p = .26; AIC^change^ = 1.3; Figure 4D). But this measure can also be meaningful on incorrect trials if we treat responses as estimation bias, i.e., how likely people are to increase or decrease the size of the chosen circles. And indeed, the estimation bias was significantly associated with VVIQ scores after accounting for the effect of change direction (*Χ*2(2) = 6.1, p = .047; AIC^change^ = −2.1; Figure 4E). In fact, the two correlation plots for correct and incorrect trials are not very different. This is confirmed by a significant overall correlation between estimation bias and VVIQ scores (r = .16, t(210) = 2.4, p = .017). We conclude that, in general, higher VVIQ scores were associated with *greater* estimation bias toward overestimating the size of the selected circles.

**Figure 4.**
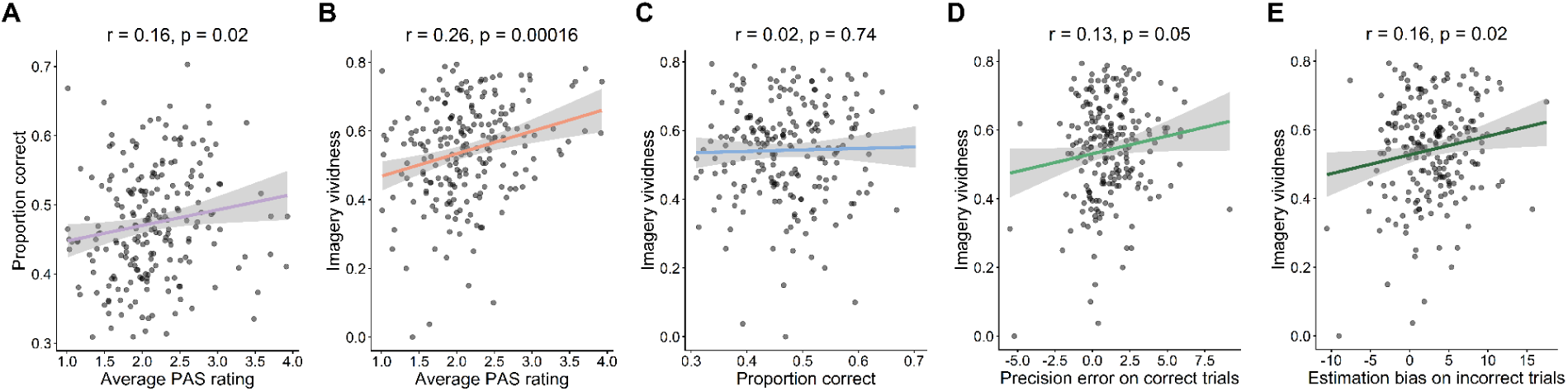
Correlations with VVIQ. **A**. The average PAS ratings and average proportions of correct trials are positively correlated. **B**. The average PAS ratings and participants total VVIQ scores are positively correlated. **C**. The average proportions of correct trials and participants total VVIQ scores are not correlated. **D**. The precision errors (averaged across correct trials) and participants total VVIQ scores are weakly positively correlated. **E**. The estimation biases (averaged across incorrect trials) and participants total VVIQ scores are positively correlated. The black dots depict individual participants in all plots. The Pearson correlation coefficients and significance levels are noted in the panel titles.

### 3.4 A closer look at incorrect trials

In the previous section, we learned that VVIQ scores correlate with estimation bias, particularly on incorrect trials. This finding aligns with the view that responses on incorrect trials are not just “blind guesses” but may well be informed by some of the memory representations related to the stimulus displays (e.g., the ‘conscious copy’), even if these representations are not veridical. Here, we ask whether the representations that guided incorrect responses may at least in part have overlapped with the representations of conscious vWM retention.

#### 3.4.1 Is estimation bias predicted by PAS ratings?

A linear mixed-effects model with the change direction of the true target as a covariate revealed a significant effect of PAS ratings (*Χ*^2^(6) = 13.3, p = .039; AIC^change^ = −1.2), with a significant interaction between PAS and change direction (*Χ*^2^(3) = 11.7, p = .008; AIC^change^ = −5.7). As can be seen from Table SI.2.5, specifically, the linear trend was significant. Figure 5A reveals that estimation bias of the wrong item exhibits the opposite effect of change direction compared to correct trials, but only if the preceding PAS rating was high. When a PAS rating of 4 was given, and the true target increased in size in the test display, but the wrong item was selected as the target, size estimation bias tended to be more often positive, i.e., the item’s size was increased rather than decreased. Conversely, when the true target decreased in size, but the wrong item was selected as the target, size estimation bias tended to be more often negative, i.e., the item’s size was decreased rather than increased (t(58) = 2.6, p = .04).

**Figure 5.**
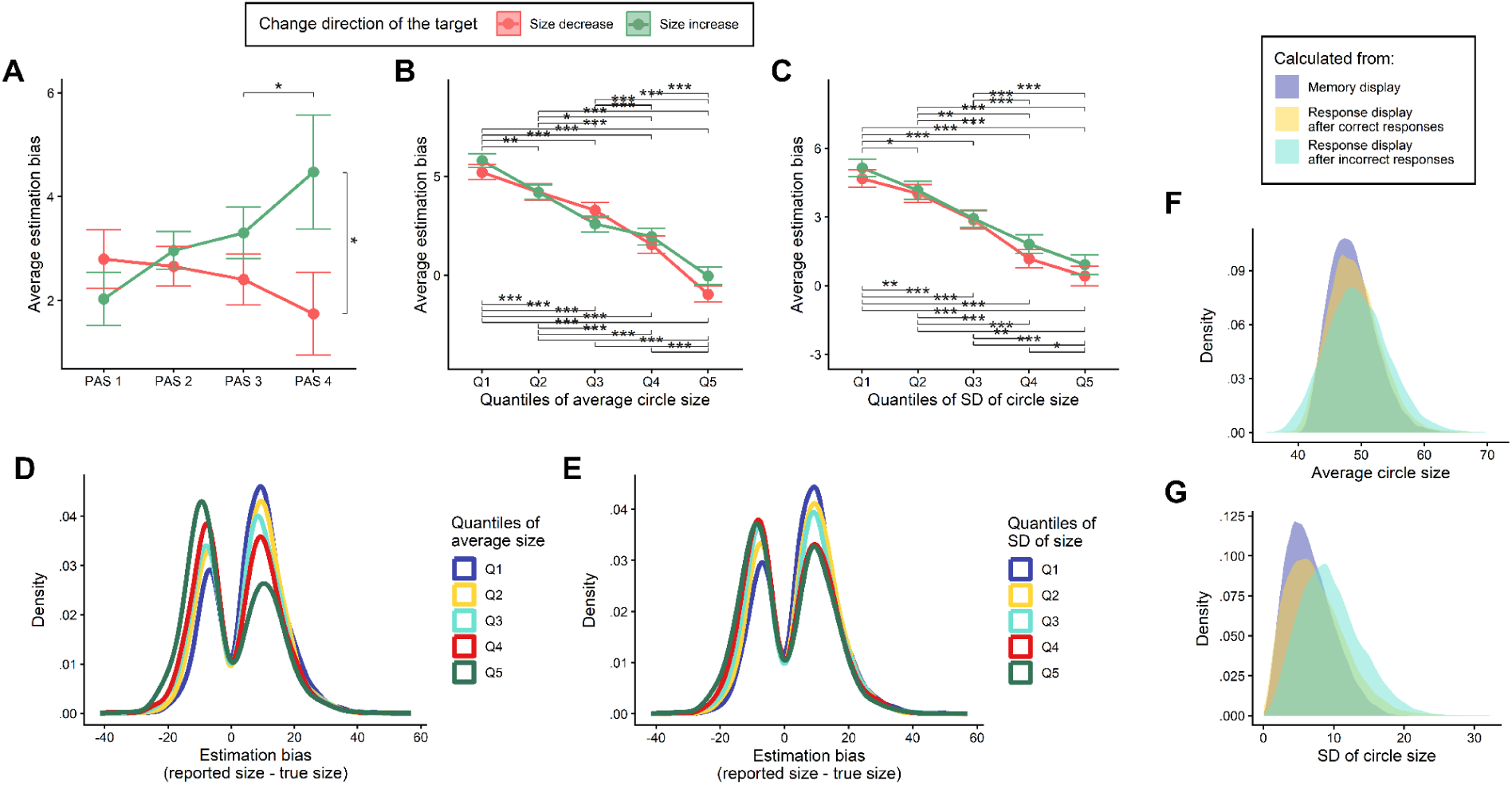
Results for incorrect trials. **A**. Estimation bias is affected by preceding (high) PAS ratings, after taking into account the change direction of the real target circle. **B**. Estimation bias is strongly affected by average circle size. **C**. Estimation bias is strongly affected by the standard deviation of circle size. **A-C** Error bars depict standard errors, and significance bars depict pairwise comparisons between factor levels with a significant t-statistic. Non-significant comparisons are omitted. **D**. Distribution of estimation bias in the entire dataset, separated by the quantiles of average circle size. **E**. Distribution of estimation bias in the entire dataset, separated by the quantiles of standard deviation of circle size. **F**. Distribution of average circle size calculated for the memory display, the final response display of correct trials, and the final response display of incorrect trials. **G**. Distribution of the standard deviation of circle size calculated for the memory display, the final response display of correct trials, and the final response display of incorrect trials.

#### 3.4.2 The effect of display statistics

Both summary statistics of the memory displays had a significant effect on the estimation bias for incorrect trials (*Χ*^2^(8) = 453.9, p < .001, AIC^change^ = −437, and *Χ*^2^(8) = 285.7, p < .001, AIC^change^ = −269, for average item size and the standard deviation of item size, respectively). The quadratic trend was significant for average item size. Figures 5B–E depict the patterns. Estimation bias was more often positive for displays with a smaller average item size and gradually became more often negative for displays with a very large average item size. The linear trend was significant for the standard deviation of item size. Estimation bias was more often positive for displays with a smaller standard deviation of item size and gradually became slightly more often negative for displays with a very large average item size. At first glance, it might be tempting to interpret this pattern as a regression to the mean. A closer inspection, however, reveals the opposite. When we calculate the same two summary statistics – average item size and the standard deviation of item size – for the final response displays, after participants had chosen a target and adjusted its size, we find that the summary statistics tend to be more extreme on incorrect trials than on correct trials (D = 0.08, p < .001 and D = 0.21, p < .001, for average item size and the standard deviation of item size, respectively). This is particularly true for the distribution of the standard deviation of item size, where the final standard deviation is often much higher than the original distribution of the metric (D = 0.29, p < .001). The distributions are depicted in Figures 5F and 5G.

Taken together with the findings that PAS ratings predict estimation bias on incorrect trials and that displays with higher standard deviation tend to yield higher PAS ratings in general, we conclude that the adjustment of circle size is guided, at least in part, by the same representations that determine participants’ trial-to-trial PAS ratings.

## 4. Discussion

The present study set out to examine how the contents of visual working memory are experienced during active maintenance and whether this subjective experience reflects the same representations that support objective task performance. Participants performed a change-detection task with continuous reporting, while additionally providing trial-by-trial ratings of their perceptual awareness of the memorized displays during the maintenance interval. Crucially, the statistical properties of the memory displays were systematically manipulated, allowing us to compare the effects of ensemble structure on both subjective and objective measures. Building on the ‘conscious copy’ model (Jacobs & Silvanto, 2015), we tested the possibility that vWM may rely on at least two distinct representations: an underlying, primarily unconscious memory trace and a consciously accessible, integrated representation of that content. If this is the case, subjective experience and objective performance should not always align and may, under certain conditions, reflect different aspects of the stored information.

### Subjective and objective measures of vWM

Considering the descriptive results of the subjective and objective measures, PAS ratings exhibited substantial variability both across and within participants, with most responses clustering around intermediate values (i.e., PAS 2), indicating that participants typically experienced only a partial or degraded representation of the memorized displays. This finding already challenges a strong version of the conscious vWM account, according to which working memory content is directly and clearly accessible to awareness (Baddeley, 2003; Skóra & Wierzchoń, 2016). Instead, the predominance of intermediate ratings suggests that conscious access to vWM content is limited and graded, consistent with previous work using subjective awareness measures (Overgaard et al., 2006).

At the same time, objective performance indicated that the task was demanding but above chance, and response precision revealed systematic biases in performance rather than random noise. Importantly, PAS ratings and correctness were weakly but significantly correlated, suggesting that subjective experience and objective performance are at least partially related. This aligns with studies linking metacognitive judgments to working memory performance (Rademaker et al., 2012; Vandenbroucke et al., 2014), but does not by itself imply that both rely on the same representation. Rather, it suggests that the representations underlying subjective experience and behavior may overlap or interact.

### Effects of ensemble properties

The most informative findings arise from the differential effects of ensemble properties on subjective and objective measures. PAS ratings increased with both average circle size and variability in circle size within the display. This pattern suggests that subjective experience is sensitive to global, ensemble-level properties, rather than being solely determined by individual item representations. Such sensitivity to summary statistics is well documented in visual perception and memory (Brady, Konkle, & Alvarez, 2011; Utochkin & Brady, 2020) and is consistent with the idea that conscious experience is inherently integrated (Bayne & Chalmers, 2003; Bayne, 2010).

In contrast, objective correctness followed a different pattern: performance was higher for a smaller average circle size and exhibited a non-linear relationship with circle-size variability. This dissociation indicates that the factors that enhance subjective clarity do not necessarily support accurate task performance. One interpretation is that while larger and more variable displays may produce a more vivid or structured conscious representation, they may simultaneously impose greater demands on the underlying memory system, reducing the fidelity of item-specific information. This pattern is difficult to reconcile with single-representation models of vWM, such as strict ‘slots’ accounts (Luck & Vogel, 1997), and instead supports the notion that different aspects of the display may be encoded or accessed differently.

Precision results further complicate this picture, as they partly align with the pattern of subjective experience and partly with the pattern of objective correctness. This intermediate pattern suggests that precision may reflect a combination of processes, potentially drawing on both item-specific and ensemble-based information. Such hybrid representations have been proposed in recent models of vWM that incorporate both discrete and continuous coding mechanisms (Brady & Tenenbaum, 2013; Orhan & Jacobs, 2014).

### Relationship between PAS and performance

The finding that PAS ratings predict correctness provides important constraints on interpretation. If subjective experience were entirely epiphenomenal, no such relationship would be expected. Instead, the observed association suggests that conscious representations can contribute to successful task performance, at least under some conditions. At the same time, the absence of a relationship between PAS and precision indicates that subjective clarity does not necessarily reflect the fine-grained fidelity of the stored information. This dissociation echoes previous findings showing that confidence judgments can be selectively related to some aspects of performance but not others (Rademaker et al., 2012).

An alternative interpretation is that PAS ratings reflect metacognitive judgments about task difficulty rather than the contents of working memory per se. Indeed, PAS has been shown to be sensitive to task-related difficulty (e.g., Skóra et al., 2021). However, this account cannot fully explain the present findings, as subjective ratings diverged from objective performance and were systematically related to estimation bias even on incorrect trials, suggesting that PAS captures aspects of the internal representations that guide behavior.

### Vividness of imagery

The relationship between VVIQ scores and task measures further supports a distinction between representations. While imagery vividness was positively associated with PAS ratings, it was unrelated to objective correctness. Instead, VVIQ scores predicted estimation bias, particularly on incorrect trials. This suggests that imagery ability primarily influences the nature of the consciously accessible representation, rather than the accuracy of the underlying memory trace.

This interpretation is consistent with evidence from aphantasia, showing that individuals with little or no visual imagery can nevertheless perform normally on visual working memory tasks (Keogh & Pearson, 2018; Bainbridge et al., 2021). Together, these findings challenge accounts that equate vWM maintenance with conscious imagery (Pearson, 2019) and instead support the idea that imagery reflects a separate representational process that can influence, but is not necessary for, task performance.

### Incorrect trials and estimation bias

The analysis of incorrect trials provides particularly strong support for the multi-representational account. Even when participants selected the wrong item, their responses exhibited systematic biases that depended on both PAS ratings and display statistics. This indicates that incorrect responses are not mere guesses, but are guided by structured internal representations.

In line with the ‘conscious copy’ model, one possibility is that when the underlying vWM representation fails to support correct identification, participants rely more strongly on a consciously accessible representation of the display. This representation may be less precise and more strongly influenced by ensemble properties, but still provides a basis for informed responding. Notably, the finding that higher PAS ratings amplify systematic biases on incorrect trials suggests that conscious representations can actively shape behavior, even when they do not correspond to the correct item.

Interestingly, the pattern of estimation bias did not reflect a simple regression to the mean, but showed an exaggeration of display-level properties. This suggests that the representations guiding responses may be constructed or transformed, rather than directly read out from memory. Such constructive processes are well documented in episodic memory (Schacter & Addis, 2007) and may also play a role in working memory when conscious representations are engaged.

### General implications

Taken together, the present findings support a view of vWM as a multi-representational system, in which subjective experience and objective performance can be partially dissociated. While both appear to draw on shared information to some extent, they differ in their sensitivity to ensemble properties and are influenced by distinct factors, such as imagery ability. This pattern is consistent with the idea that vWM involves both an underlying, largely unconscious representation and a consciously accessible, integrated representation, as proposed by the ‘conscious copy’ model (Jacobs & Silvanto, 2015). Crucially, our results go beyond a simple coexistence account and suggest that these representations may differ not only in their accessibility, but also in their format: whereas the underlying representation may preserve item-specific information (while also being sensitive to ensemble-level properties to some extent), the consciously accessible representation appears to be more strongly shaped by global, ensemble-level properties of the display. This distinction provides a potential resolution to the long-standing tension between ‘slots’-based models, which assume independent item storage (Luck & Vogel, 1997), and evidence for structured, ensemble-based representations in vWM (Brady et al., 2011; Utochkin & Brady, 2020).

More broadly, these results suggest that trial-by-trial performance in vWM tasks may not always reflect the same underlying processes. Instead, participants may flexibly rely on different representations depending on task demands and the quality of the available information. In situations where the underlying memory trace is sufficiently strong, behavior may be primarily driven by item-specific representations. However, when this representation is degraded or insufficient, participants may increasingly rely on a consciously accessible, more global representation of the display. This interpretation is consistent with the idea that cognitive systems adaptively combine multiple sources of information to optimize performance under uncertainty (Brady & Tenenbaum, 2013; Orhan & Jacobs, 2014). Importantly, it also implies that variability in behavioral responses across trials may reflect shifts in representational strategy, rather than noise alone.

These considerations have important implications for both theoretical models and the interpretation of neural data (e.g., Oh, Kim & Kang, 2019). If multiple representational formats can contribute to performance, then neural signals associated with vWM may reflect a mixture of processes, including both activity related to latent memory traces and activity associated with consciously accessible representations (Stokes, 2015; Trübutschek et al., 2017). As a result, averaging across trials without accounting for differences in subjective experience or strategy may obscure meaningful distinctions in the underlying neural mechanisms. Incorporating trial-by-trial subjective measures, such as PAS ratings, may therefore provide a valuable tool for disentangling these contributions.

### Limitations and future directions

Several limitations should be noted. First, the interpretation of PAS ratings as reflecting conscious experience depends on participants’ understanding and use of the scale. Although participants were carefully instructed and PAS has been widely used as a measure of perceptual awareness (Ramsøy & Overgaard, 2004), alternative interpretations in terms of metacognitive judgments or task-related heuristics cannot be fully ruled out. Future studies could address this issue by combining PAS with additional measures, such as confidence ratings or objective awareness indices, to more precisely characterize the nature of the reported experience.

Second, the present study provides behavioral evidence for multiple representations, but does not directly identify their neural correlates. While the observed dissociations are consistent with proposals of distinct functional states of working memory, including ‘activity-silent’ representations (Stokes, 2015; Trübutschek et al., 2017), direct evidence for such mechanisms in the present paradigm is lacking. Combining the current behavioral approach with neuroimaging or electrophysiological methods would allow for a more direct test of whether the proposed representations correspond to distinct neural processes.

Third, although the manipulation of ensemble properties proved informative, the present design does not fully disentangle their contribution from other stimulus dimensions, and the correlation between average size and variability limits the extent to which their effects can be interpreted independently. Future work could employ more orthogonal manipulations or computational modeling approaches to better isolate the mechanisms underlying these effects.

Finally, further research is needed to establish the generality of the present findings across different task paradigms, stimulus types, and temporal dynamics of working memory. In particular, it remains an open question whether similar dissociations between subjective experience and objective performance would be observed in tasks that rely less on spatially distributed displays or that place different demands on memory precision and decision-making.

## Supporting information

Supplementary Information for the manuscript

## Acknowledgements

We thank Kristian Sandberg and the entire NeuralArchCon consortium for their concerted effort (www.neuralarchcon.org). We thank Tomasz Kostka, Wiktoria Jakubowska, and the C-lab interns from 2020 to 2022 for their assistance with behavioral data collection. We thank Justyna Hobot and Krystian Barzykowski for translating the VVIQ, and Timo Kvamme for preparing the VVIQ data for analysis.

## Data Availability Statement

The scripts used for data preprocessing and statistical analysis, together with raw data and a CSV file with preprocessed data, are available for download from the OSF repository (https://osf.io/q8hpt/).

## Author Contributions

**AL:** data curation, formal analysis, investigation, software, visualization. **KC:** investigation, writing – original draft. **RR:** conceptualization, data curation, formal analysis, funding acquisition, investigation, methodology, project administration, software, supervision, visualization, writing – original draft.

