## Supplementary Information for the manuscript for "The Two Lives of Visual Working Memory: Evidence for Distinct Conscious and Unconscious Representations"

### SI.1 Examples of memory displays

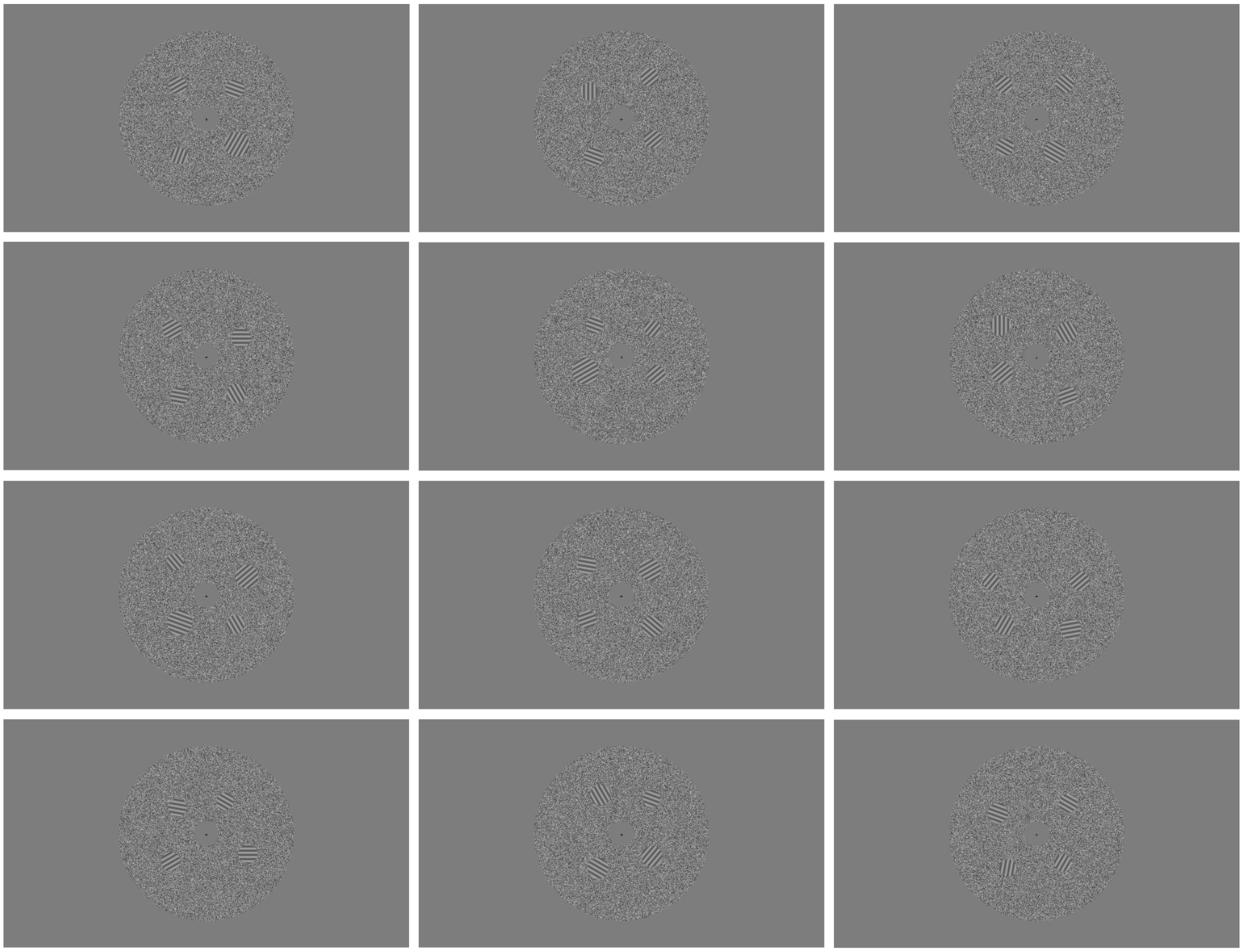

**Figure SI.1.** The memory displays consisted of an ensemble of four circles, placed semi-randomly in the four quadrants of the noise annulus.

### SI.2 Model fits

**Table SI.2.1.** Model fits for PAS (corresponding to Results section 3.2.1)

| **Model (clmm):** | **PAS ~ average circle size + (1\|ID)** | | | |
|---|---|---|---|---|
| AIC change: | 67799.46 - 68073.05 = -273.59 | | | |
| Fixed effects: | Estimate | SE | z | probability |
| Linear | 0.39 | 0.02 | 16.35 | < 2e-16 *** |
| Quadratic | 0.1 | 0.02 | 4.29 | 1.79e-05 *** |
| Cubic | -0.003 | 0.02 | -0.13 | 0.9 |
| Random effect: | Variance | SD | | |
| | 4.09 | 2.02 | | |
| **Model (clmm):** | **PAS ~ SD of circle size + (1\|ID)** | | | |
| AIC change: | 67601.82 - 68073.05 = -471.23 | | | |
| Fixed Effects: | Estimate | SE | z | probability |
| Linear | 0.47 | 0.02 | 19.9 | <2e-16 *** |
| Quadratic | 0.22 | 0.02 | 9.41 | <2e-16 *** |
| Cubic | 0.0004 | 0.02 | 0.02 | 0.98 |
| Random effect: | Variance | SD | | |
| | 4.12 | 2.03 | | |

**Table SI.2.2.** Model fits for %correct (corresponding to Results sections 3.1.4 and 3.2.2)

| **Model (glmer):** | **%correct ~ PAS + (1\|ID)** | | | |
|---|---|---|---|---|
| AIC change: | 3652.5 - 3679.5 = -27 | | | |
| Fixed effects: | Estimate | SE | z | probability |
| (Intercept) | -0.07 | 0.024 | -2.89 | 0.0039 ** |
| Linear | 0.2 | 0.041 | 4.84 | 1.32e-06 *** |
| Quadratic | 0.001 | 0.032 | 0.04 | 0.97 |
| Cubic | -0.01 | 0.022 | -0.52 | 0.6 |
| Random effect: | Variance | SD | | |
| | 0.08 | 0.28 | | |
| **Model (glmer):** | **%correct ~ average circle size + (1\|ID)** | | | |
| AIC change: | 5839.5 - 5854.6 = -15.1 | | | |

| Fixed effects: | Estimate | SE | z | probability |
|---|---|---|---|---|
| (Intercept) | -0.11 | 0.022 | -4.94 | 7.76e-07 *** |
| Linear | -0.08 | 0.023 | -3.26 | 0.0011 ** |
| Quadratic | 0.01 | 0.023 | 0.61 | 0.54 |
| Cubic | -0.03 | 0.023 | -1.17 | 0.24 |
| Random effect: | Variance | SD | | |
| | 0.08 | 0.29 | | |
| **Model (glmer):** | **%correct ~ SD of circle size + (1\|ID)** | | | |
| AIC change: | 5825.4 - 5873.9 = -48.5 | | | |
| Fixed Effects: | Estimate | SE | z | probability |
| (Intercept) | -0.11 | 0.022 | -4.89 | 1.00e-06 *** |
| Linear | -0.02 | 0.023 | -0.97 | 0.33 |
| Quadratic | 0.15 | 0.023 | 6.61 | 3.83e-11 *** |
| Cubic | -0.07 | 0.023 | -3.16 | 0.0016 ** |
| Random effect: | Variance | SD | | |
| | 0.08 | 0.29 | | |

**Table SI.2.3.** Model fits for precision (correct trials only; corresponding to Results sections 3.1.4 and 3.2.3)

| **Model (glmer):** | **precision ~ PAS * change direction + (1\|ID)** | | | |
|---|---|---|---|---|
| AIC change: | 7253.3 - 7246.0 = 7.3 | | | |
| Fixed effects: | Estimate | SE | t | probability |
| (Intercept) | 3.97 | 0.17 | 23.05 | <2e-16 *** |
| Linear | 0.13 | 0.38 | 0.35 | 0.73 |
| Quadratic | -0.29 | 0.34 | -0.85 | 0.39 |
| Cubic | -0.22 | 0.3 | -0.74 | 0.46 |
| change direction | -6.34 | 0.24 | -26.08 | <2e-16 *** |
| Linear * change dir. | 0.39 | 0.54 | 0.73 | 0.46 |
| Quadratic * ch.dir. | 0.09 | 0.49 | 0.19 | 0.85 |
| Cubic * change dir. | 0.17 | 0.43 | 0.39 | 0.7 |
| Random effect: | Variance | SD | | |
| | 9.6e-15 | 9.8e-08 | | |

| **Model (glmer):** | **precision ~ average circle size * change direction + (1\|ID)** | | | |
|---|---|---|---|---|
| AIC change: | 11766 - 11773 = -7 | | | |
| Fixed effects: | Estimate | SE | t | probability |
| (Intercept) | 4.07 | 0.12 | 33.28 | <2e-16 *** |
| Linear | -0.25 | 0.25 | -1.01 | 0.31 |
| Quadratic | -0.49 | 0.25 | -1.96 | 0.051 . |
| Cubic | -0.06 | 0.25 | -0.26 | 0.8 |
| change direction | -6.27 | 0.158 | -39.8 | <2e-16 *** |
| Linear * change dir. | -0.79 | 0.35 | -2.25 | 0.02 * |
| Quadratic * ch.dir. | 0.57 | 0.35 | 1.63 | 0.1 |
| Cubic * change dir. | -0.02 | 0.35 | -0.07 | 0.95 |
| Random effect: | Variance | SD | | |
| | 0.54 | 0.73 | | |
| **Model (glmer):** | **precision ~ SD of circle size * change direction + (1\|ID)** | | | |
| AIC change: | 11722 - 11745 = -23 | | | |
| Fixed Effects: | Estimate | SE | t | probability |
| (Intercept) | 4.06 | 0.12 | 33.1 | < 2e-16 *** |
| Linear | 0.52 | 0.25 | 2.08 | 0.038 * |
| Quadratic | -0.85 | 0.25 | -3.44 | 0.0006 *** |
| Cubic | -0.07 | 0.25 | -0.29 | 0.77 |
| change direction | -6.28 | 0.16 | -39.95 | < 2e-16 *** |
| Linear * change dir. | -1.54 | 0.35 | -4.39 | 1.2e-05 *** |
| Quadratic * ch.dir. | 1.14 | 0.35 | 3.26 | 0.001 ** |
| Cubic * change dir. | -0.37 | 0.35 | -1.05 | 0.29 |
| Random effect: | Variance | SD | | |
| | 0.6 | 0.77 | | |

**Table SI.2.4.** Model fits for VVIQ (corresponding to Results section 3.3)

| **Model (glmer):** | **estimation bias (correct trials only) ~ PAS * change direction + (1\|ID)** | | | |
|---|---|---|---|---|
| AIC change: | 2158.2 - 2156.9 = 1.3 | | | |
| Fixed effects: | Estimate | SE | t | probability |
| (Intercept) | 3.7 | 0.82 | 4.51 | 8.3e-06 *** |
| VVIQ | 0.66 | 1.46 | 0.45 | 0.65 |

| | | | | |
|---|---|---|---|---|
| change direction | -7.16 | 1.16 | -6.17 | 1.6e-09 *** |
| VVIQ * change dir. | 1.65 | 2.07 | 0.8 | 0.43 |
| Random effect: | Variance | SD | | |
| | 0 | 0 | | |
| **Model (glmer):** | **estimation bias (incorrect trials only) ~ PAS * change direction + (1\|ID)** | | | |
| AIC change: | 2287.7 - 2289.8 = -2.1 | | | |
| Fixed effects: | Estimate | SE | t | probability |
| (Intercept) | -0.2 | 1.2 | -0.17 | 0.87 |
| VVIQ | 5.26 | 2.13 | 2.47 | 0.014 * |
| change direction | 1.12 | 0.81 | 1.38 | 0.17 |
| VVIQ * change dir. | -1.44 | 1.44 | -0.99 | 0.32 |
| Random effect: | Variance | SD | | |
| | 15.3 | 3.92 | | |

**Table SI.2.5.** Model fits for estimation bias on incorrect trials (corresponding to Results section 3.4)

| **Model (glmer):** | **estimation bias ~ PAS * change direction + (1\|ID)** | | | |
|---|---|---|---|---|
| AIC change: | 8253.6 - 8254.8 = -1.2 | | | |
| Fixed effects: | Estimate | SE | t | probability |
| (Intercept) | 2.32 | 0.36 | 6.51 | 2.5e-10 *** |
| Linear | -1.01 | 0.52 | -1.94 | 0.053 . |
| Quadratic | -0.39 | 0.46 | -0.85 | 0.4 |
| Cubic | -0.13 | 0.4 | -0.33 | 0.74 |
| change direction | 0.68 | 0.33 | 2.07 | 0.038 * |
| Linear * change dir. | 2.42 | 0.73 | 3.31 | 0.001 *** |
| Quadratic * ch.dir. | 0.24 | 0.65 | 0.37 | 0.71 |
| Cubic * change dir. | 0.49 | 0.57 | 0.86 | 0.39 |
| Random effect: | Variance | SD | | |
| | 15.7 | 3.97 | | |
| **Model (glmer):** | **estimation bias ~ average circle size * change direction + (1\|ID)** | | | |
| AIC change: | 12933 - 13370 = -437 | | | |
| Fixed effects: | Estimate | SE | t | probability |

| | | | | |
|---|---|---|---|---|
| (Intercept) | 2.65 | 0.3 | 8.71 | 3.7e-16 *** |
| Linear | -4.74 | 0.29 | -16.24 | < 2e-16 *** |
| Quadratic | -1.02 | 0.29 | -3.51 | 0.0005 *** |
| Cubic | -0.27 | 0.29 | -0.94 | 0.35 |
| change direction | 0.25 | 0.18 | 1.33 | 0.18 |
| Linear * change dir. | 0.34 | 0.41 | 0.82 | 0.41 |
| Quadratic * ch.dir. | 1.07 | 0.41 | 2.6 | 0.009 ** |
| Cubic * change dir. | -0.16 | 0.41 | -0.38 | 0.71 |
| Random effect: | Variance | SD | | |
| | 16.4 | 4.04 | | |
| **Model (glmer):** | **estimation bias ~ SD of circle size * change direction + (1\|ID)** | | | |
| AIC change: | 12836 - 13105 = -269 | | | |
| Fixed Effects: | Estimate | SE | t | probability |
| (Intercept) | 2.64 | 0.31 | 8.62 | 7.1e-16 *** |
| Linear | -3.59 | 0.28 | -12.6 | < 2e-16 *** |
| Quadratic | -0.2 | 0.28 | -0.7 | 0.49 |
| Cubic | 0.46 | 0.28 | 1.61 | 0.11 |
| change direction | 0.36 | 0.18 | 1.99 | 0.046 * |
| Linear * change dir. | 0.18 | 0.4 | 0.44 | 0.66 |
| Quadratic * ch.dir. | 0.27 | 0.4 | 0.68 | 0.5 |
| Cubic * change dir. | -0.31 | 0.4 | -0.77 | 0.44 |
| Random effect: | Variance | SD | | |
| | 16.7 | 4.09 | | |

### SI.3 The effects of target properties on working memory performance

Three target properties were manipulated in this experiment: target location (the target is in one of the four visual field quadrants), relative target size (the target is either the smallest, the second smallest, the largest, or the second largest item in the stimulus array), and size change direction (the target reduced or increased in size compared to its original radius).

#### *SI.3.1 Correctness*

Relative target size yielded the largest test statistic ($X^2(3)=991.5$, $p<.001$, $AIC^{change} = -985.4$). Participants were much more likely to correctly identify the target if it was the largest item in the array (see Figure SI.3.1.A). The second largest item also showed improved correctness compared to the two smaller array items, while still yielding systematically lower correctness than the largest item. Change direction did not have a systematic effect on correctness ($X^2(1)=2.9$, $p=.09$, $AIC^{change} = -0.9$), even though a slight trend towards better detection of targets that reduced in size can be discerned from Figure SI.3.1.B.

The effect of target location deserves some closer attention. As can be seen in Figure SI.3.1.C, participants were more likely to correctly identify the target when it appeared in the upper visual field, i.e., in the upper right or upper left quadrant ($X^2(3)=155.8$, $p<.001$, $AIC^{change} = -149.8$). Performance was worst in the lower right quadrant and somewhat improved in the lower left quadrant, but remained clearly inferior to performance in the upper visual field locations. It should be noted, however, that participants also showed a clear bias toward selecting the upper visual field items as targets, even when that choice was incorrect (see section SI.3.4). It is therefore likely that some of the effect of target location is carried by this selection bias.

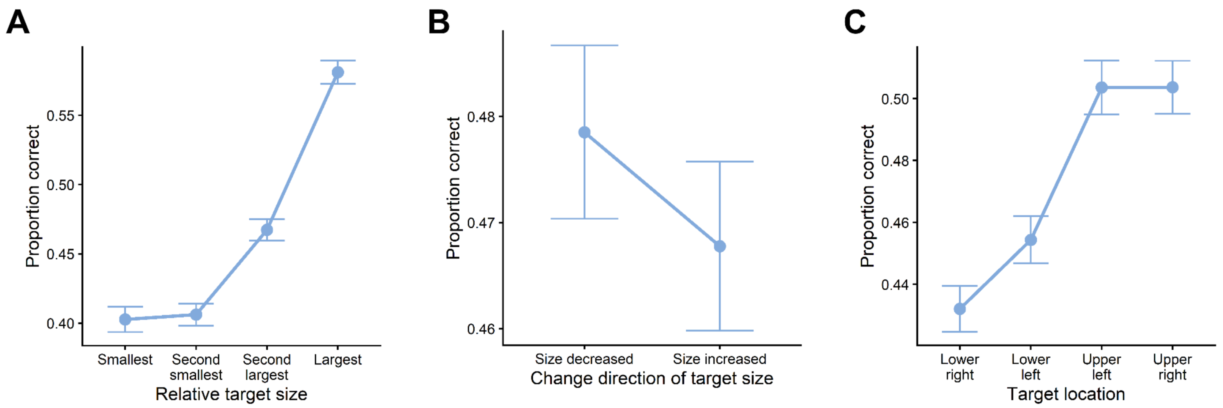

**Figure SI.3.1.** Effect of target properties on the proportion of correct trials. **A**. Effect of relative target size. **B**. Change direction. **C**. Target location. Error bars depict standard errors.

### *SI.3.2 Precision*

All three target properties that were manipulated in the paradigm (target location, relative target size, and size change direction) had a significant effect on the precision of target adjustment ($X^2(3) = 28.8$, p = .001, $AIC^{change} = -22.8$; $X^2(1) = 313.9$, p < .001, $AIC^{change} = -311.9$, and $X^2(3) = 16.2$, p < .001, $AIC^{change} = -10.2$, for relative target size, change direction, and target location, respectively). The precision error tended to be lowest if the target was the second-largest item in the array. Precision did not differ systematically when the target was one of the smaller items or the largest item in the array (all t<1.8, all p>.08). As noted above, change direction influenced the precision of responses the most. When the target increased in size, participants tended to overestimate the original target size by an average of 4 pixels. When the target decreased in size, participants tended to underestimate its original size by an average of 2.2 pixels. Finally, precision tended to be higher in the upper visual field compared to the lower visual field.

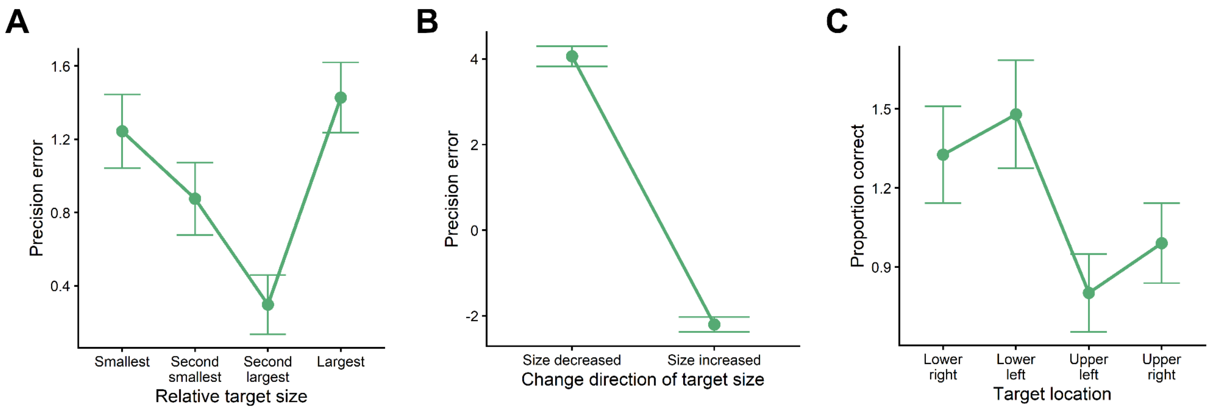


**Figure SI.3.2.** Effect of target properties on the precision of the size adjustment of the target. **A**. Effect of relative target size. **B**. Change direction. **C**. Target location. Error bars depict standard errors.

*SI.3.3 PAS*

PAS ratings were given before the test display; therefore, properties of the target item should not have influenced PAS ratings. This was indeed the case. None of the regression models with target location, relative target size, or size-change direction as predictors yielded statistically significant results (test statistics not reported).

Since objective vWM, particularly correctness, was best for relatively larger targets in the upper visual field, however, we wished to assess whether similar patterns can also be discerned from PAS ratings. To answer this question, we calculated the size difference of the average of the two upper items and the average of the two lower items on each trial, and again divided these differences into quantiles. The same was done for the size difference between the two left items and the two right items, leading to a comparison of the upper vs. lower visual fields and the left vs. right visual fields.

Both visual field asymmetries affected PAS ratings, with a U-shaped pattern in both cases (see Figure SI.3.3; $X^2(3) = 132.9$, $p < .001$, $AIC^{change} = -124.9$, and $X^2(3) = 134.6$, $p < .001$, $AIC^{change} = -126.6$, for the up/down and left/right quantiles, respectively). PAS ratings were highest when there was a large asymmetry of item sizes between the upper/lower and left/right visual fields, regardless of the direction of this asymmetry. Pairwise comparisons of the up/down asymmetry quantiles confirmed that the quantiles -2 and 2 do not differ from each other ($t(216)=0.6$), but differ systematically from the quantiles -1, 0, and 1 (all $t(216)>5.4$, $p<0.001$). The comparison between the left and right visual fields resulted in largely the same pattern. The quadrants -2 and 2 tend to yield the highest PAS ratings and are not different from each other ($t(216) =- 0.6$), but both differ significantly from quadrants -1, 0, and 1 (all $t(216) > 5.8$, $p < 0.001$). None of the quadrants (-1, 0, and 1) yield systematically different PAS ratings (all $t(216) < 1.5$, $p > .94$).

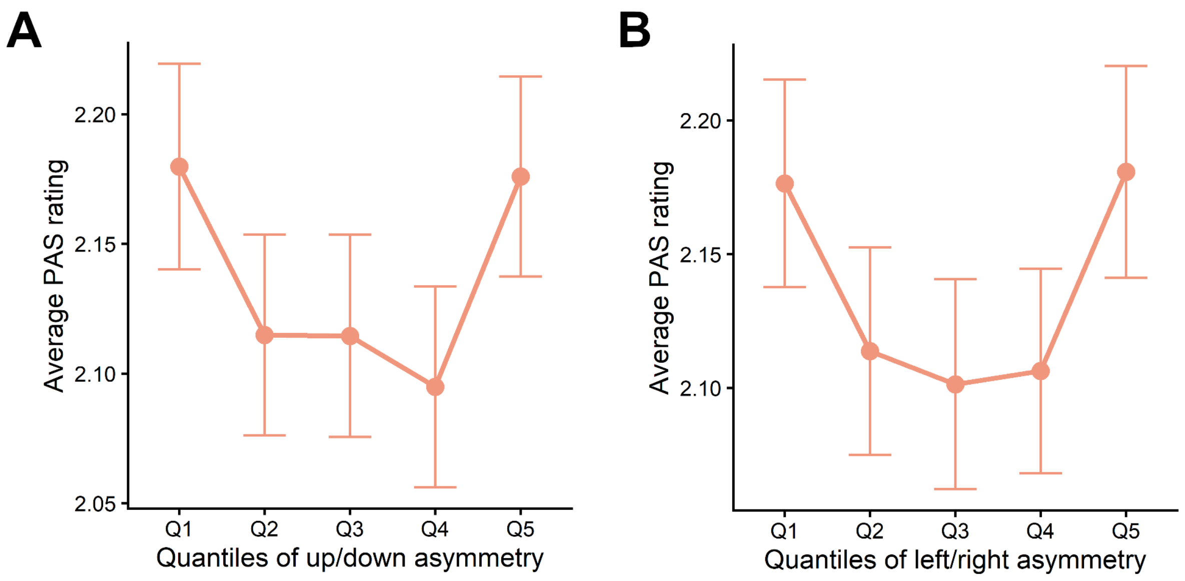


**Figure SI.3.3.** Effect of visual field asymmetries on PAS ratings. **A**. Effect of the asymmetry between the upper and the lower visual field. **B**. Effect of the asymmetry between the left and the right visual field. Error bars depict standard errors.

*SI.3.4 Incorrect trials*

We next asked which item properties made them more likely to be incorrectly selected. We call this selection bias. Item location had a significant effect on selection bias ($X^2(6)=87.1$, $p<.001$, $AIC^{change} = -75.1$), with the items in the upper right and left locations being somewhat more likely to be selected than the items in the lower visual field (Figure SI.3.4.A). This selection pattern is quite similar to the one observed on correct trials, but the interaction between correctness and item location is nevertheless significant ($X^2(3)=8.6$, $p=.035$, $AIC^{change} = -2.6$). The lower-right location is more likely to be selected on incorrect trials than on correct trials, and the upper-left location is less likely to be selected on incorrect trials than on correct trials.

Looking at the estimation bias, item location had a significant effect on that metric as well ($X^2(3)=14.2$, $p = .0026$, $AIC^{change} = -8.3$). Estimation bias tended to be slightly more positive in the upper visual field (see Figure SI.3.4.C).

Instead of the relative item size in the *memory* display, we analysed the relative item size in the *test* display for selection bias. This factor had a pronounced effect on the selection bias in the incorrect trials ($X^2(6)=1750$, $p<.001$, $AIC^{change} = -1737.7$), with larger items being increasingly likely to be selected (Figure SI.3.4.B). This selection pattern is very different from the one observed on correct trials (interaction with correctness $X^2(3)=1010.7$, $p<.001$, $AIC^{change} = -1004.7$), where the largest and the smallest items are much more likely to be selected than the intermediary ones.

Looking at the estimation bias, relative item size had a significant effect on that metric as well ($X^2(3)=55$, $p<.001$, $AIC^{change} = -53$). Estimation bias tended to be positive when the smallest item was selected and gradually became negative when one of the larger items was selected.

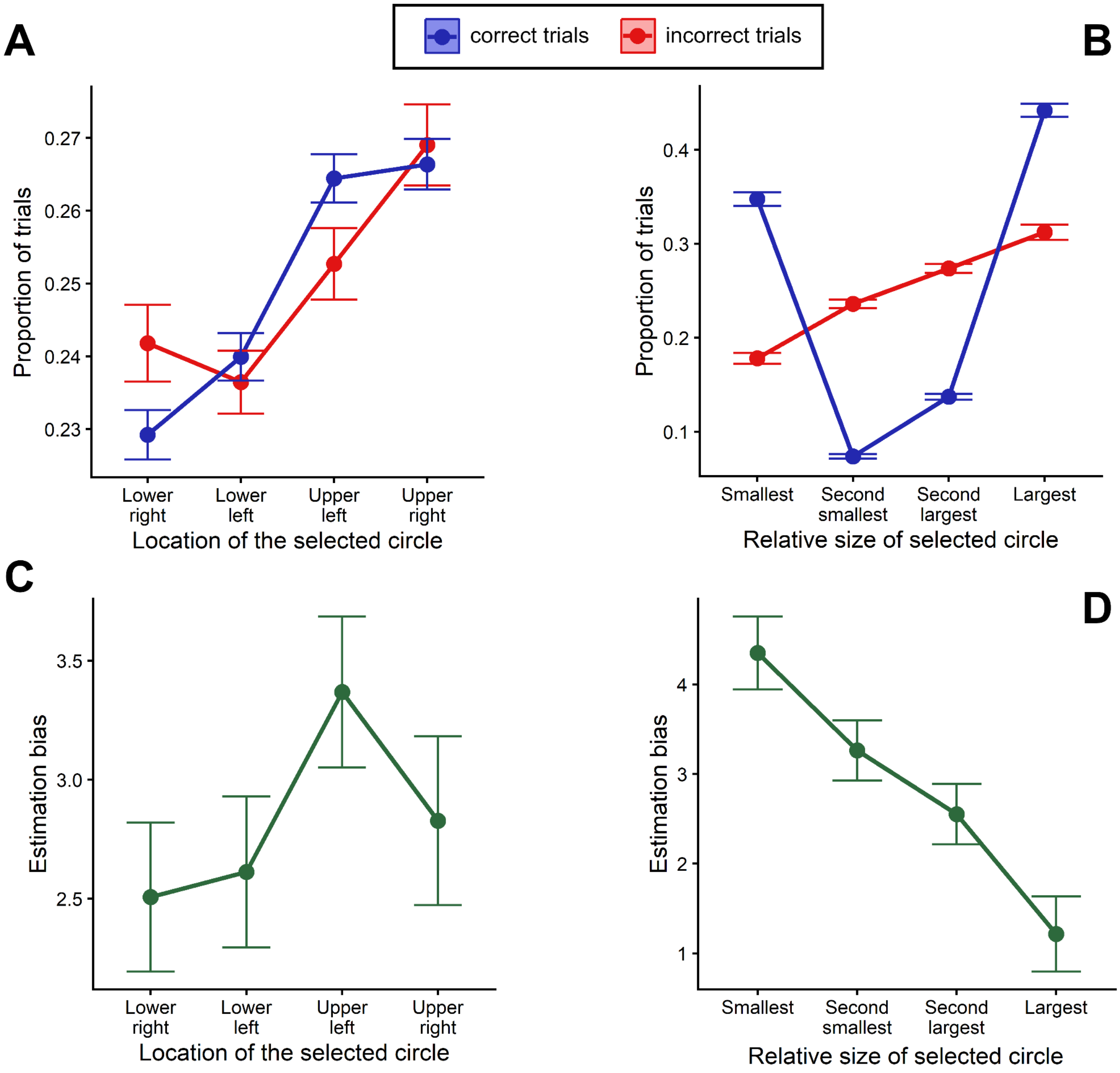


**Figure SI.3.4.** Circle selection probability/estimation bias depending on location and size. **A.** Proportion of trials where each of the four locations in the test display was selected, separately for correct and incorrect trials. **B**. Proportion of trials where circles of relative sizes were selected in the test display, separately for correct and incorrect trials. **C**. Average estimation bias on incorrect trials for each of the four locations. **D**. Average estimation bias on incorrect trials for relative circle sizes. Error bars depict standard errors.
